# Allosteric binders of ACE2 are promising anti-SARS-CoV-2 agents

**DOI:** 10.1101/2022.03.15.484484

**Authors:** Joshua E. Hochuli, Sankalp Jain, Cleber Melo-Filho, Zoe L. Sessions, Tesia Bobrowski, Jun Choe, Johnny Zheng, Richard Eastman, Daniel C. Talley, Ganesha Rai, Anton Simeonov, Alexander Tropsha, Eugene N. Muratov, Bolormaa Baljinnyam, Alexey V. Zakharov

## Abstract

The COVID-19 pandemic has had enormous health, economic, and social consequences. Vaccines have been successful in reducing rates of infection and hospitalization, but there is still a need for an acute treatment for the disease. We investigate whether compounds that bind the human ACE2 protein can interrupt SARS-CoV-2 replication without damaging ACE2’s natural enzymatic function. Initial compounds were screened for binding to ACE2 but little interruption of ACE2 enzymatic activity. This set of compounds was extended by application of quantitative structure-activity analysis, which resulted in 512 virtual hits for further confirmatory screening. A subsequent SARS-CoV-2 replication assay revealed that five of these compounds inhibit SARS-CoV-2 replication in human cells. Further effort is required to completely determine the antiviral mechanism of these compounds, but they serve as a strong starting point for both development of acute treatments for COVID-19 and research into the mechanism of infection.

TOC Graphic: Overall study design.

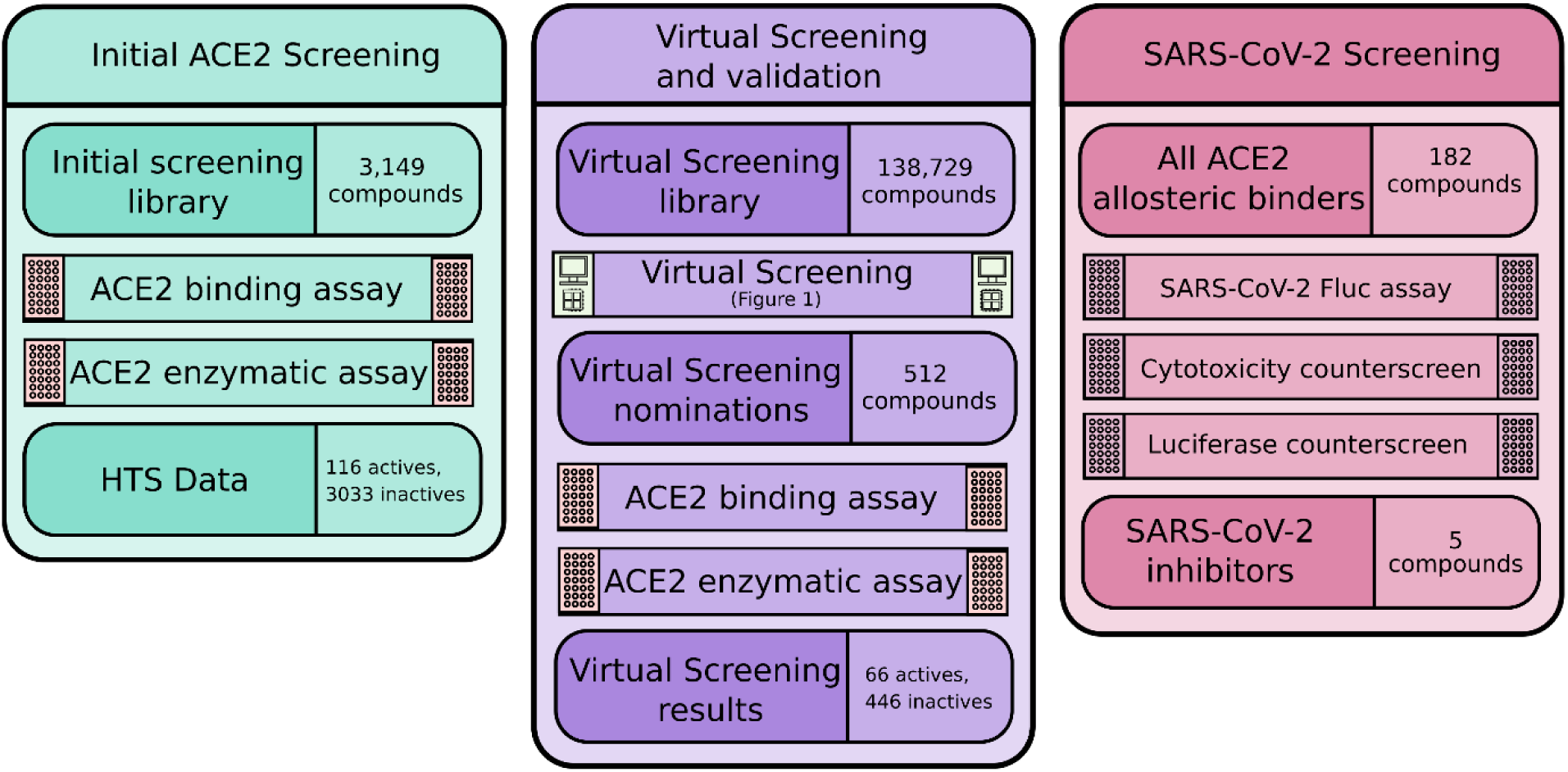

## Introduction

The radical consequences of coronavirus disease 2019 (COVID-19) and the virus that causes it, severe acute respiratory syndrome coronavirus 2 (SARS-CoV-2), since its advent in late 2019, are evident in many facets of daily life. Since this time, many economies around the world have entered recessions, unemployment rates have dramatically increased, and the healthcare infrastructure in many countries is being pushed to its limits.^1–3^ More significantly, COVID-19 has killed almost 6 million people worldwide and has infected well over 400 million as of February 2022.^4^

Vaccine deployment has been a general success in reducing rates of infection^5^ and especially rates of hospitalization^5,6^, though less so for transmission from a vaccinated individual.^7^ Importantly, there are breakthrough cases where vaccinated individuals contract the virus, which can still result in severe symptoms or death. Variants of the virus are actively developing and spreading^8^, and there is always a possibility that a viral mutation will allow evasion of the immune system detection provided by a previous infection or vaccine. Thus, while the effort that the scientific community has put into developing a vaccine is incredibly important, it is still necessary to dedicate effort to finding therapies for acute COVID-19 to alleviate the severity of infection.

It is imperative that efforts be dedicated to the development of small molecule inhibitors targeting various parts of the viral life cycle, either by direct action upon viral proteins or upon host factors that play integral roles in viral replication. These therapies would not only be immediately applicable to COVID-19, but also useful in any future coronavirus epidemics.^9^ Thus far, there are several repurposed drugs granted emergency use authorization in the treatment of COVID-19. Remdesivir, a nucleoside analog that inhibits the action of SARS-CoV-2 RNA polymerase II, has taken a premier role in treatment since early 2020. Other drugs that are currently being used to treat COVID-19 cases are the rheumatoid arthritis drug baricitinib and the corticosteroid dexamethasone, which act to reduce the inflammation associated with severe infection.^10^ There is also at least one approved new drug for treatment of COVID-19^11^, but its efficacy in practice remains to be seen.

Like other research organizations, The National Center for Advancing Translational Sciences (NCATS) has initiated efforts to support and promote the development of anti-SARS-CoV-2 treatments. The OpenData Portal for its COVID-19 drug repurposing^12^ allows researchers and public health officials to expedite the development of SARS-CoV-2 interventions through open data sharing and analysis tools, and to prioritize antiviral discovery for further development in treating COVID-19. Furthermore, a number of researchers have been performing large-scale virtual screening of the in-house libraries with the aim to identify chemotypes with antiviral activity and limited host cell cytotoxicity.^13,14^ Also, various screening assays have been set up that allow HTS testing to speed up the experimental process.^15–17^

Although many small molecule therapies for COVID-19 thus far have targeted viral proteins, there is interest in discovering small molecule inhibitors of host proteins integral to the SARS-CoV-2 life cycle. Developing drugs to target important host proteins reduces the chance of the virus developing drug resistance.^18^ One such host protein of particular interest is the primary entry receptor for SARS-CoV-2, angiotensin converting enzyme 2, or ACE2.^19^

ACE2 is a key enzyme in the Renin-Angiotensinogen-Aldosterone System (RAAS), which is implicated in renal, pulmonary, immune, and cardiovascular function.^20–22^ ACE2 is responsible for the conversion of angiotensin II to ANG 1-9, and by further extension anti-inflammatory effects.^23^ ANG I is cleaved into angiotensin II (ANG II) by angiotensin converting enzyme (ACE). ANG II is responsible for binding to AT1R and signaling prostaglandin and aldosterone production, increasing Na+ retention, blood pressure, and inflammation. ANG II is converted into angiotensin III (ANG 1-7) by ACE2, maintaining homeostasis by binding to MAS-R, producing anti-inflammatory, anti-fibrotic effects.^24,25^ This signaling is completed using soluble ACE2, which is shed from its cellular state by disintegrin and metalloproteinase 17 (ADAM17) as well as TMPRSS22625.

Due to the complex role of the RAAS system, the effects of inhibiting ACE2 enzymatic activity include increased inflammation, fibrosis, oxidative stress, and vasoconstriction.^25^ In the event of ACE2 loss-of-function due to SARS-CoV-2 binding or an orthosteric ACE2 inhibitor^27^, downstream reduction in production of ANG (1-7) would hinder anti-inflammatory compensation as well as allow for the over production and expression of ANG II, inciting further inflammation and fibrosis. Therefore, inhibition of the SARS-CoV-2 spike protein interaction without disturbing natural enzymatic activity should be the focus of ACE2 inhibition^28^ as it applies to COVID-19 therapeutics.

Understanding the powerful abilities of computational tools when used correctly^29,30^ enabled many research groups to find COVID-19 therapeutics as well as to prevent the binding of SARS-CoV-2 to ACE2^31^. In some cases, molecular modeling has been used to virtually screen compounds^32^ and identify predicted synergistic mixtures^33^ that may aid in the treatment of COVID-19. We report here a hybrid discovery approach, where computational models are in conjunction with highthroughput screening to establish as large a dataset as possible of compounds with desired properties.

The aims of this study are (i) to identify allosteric binders of ACE2 without enzyme inhibitory activity and (ii) to discover small molecules acting under this mechanism as potential agents against the SARS-CoV-2 virus. To do so, we utilized assays for ACE2 binding and enzymatic activity to screen a large dataset of small molecules and develop initial data for modeling. Independent modeling efforts were undertaken by both NCATS and UNC teams to nominate a diverse set of molecules predicted to bind to ACE2 without significant interruption of enzymatic activity. Computationally nominated compounds were experimentally screened, resulting in 68 actives (out of 512 nominations; hit rate ∼13%). The confirmed hits were then subjected to an assay for inhibitions of SARS-CoV-2 replication in human cells. Appropriate counterscreens for cytotoxicity and luciferase assay interference were applied to all hits. In total, five compounds were identified as inhibitors of SARS-CoV-2 replication at the micromolar level presumably due to ACE2 binding without significant inhibition of natural ACE2 enzymatic activity.

## Results and Discussion

### Initial Screening

To identify small molecule binders to ACE2 which does not interfere with its enzymatic activity, we utilized microscale thermophoresis (MST) in combination with an enzymatic assay. Extended assay descriptions are included in the Supplementary Methods.

Recombinant polyhistidine-tagged (His-tag) extracellular domain of ACE2 and the fluorogenic substrate MCA ((7-Methoxycoumarin-4-acetic acid) - Ala - Pro – Lys (Dnp) – OH) was used for the enzymatic assay. For MST, the recombinant ACE2 protein was labeled with a His-tag specific fluorophore to monitor any binding events. MLN-4760, a known ACE2 inhibitor, was used to test and optimize the conditions of the enzymatic and MST assays. In the enzymatic assay MLN-4760 showed a dose-dependent inhibition of ACE2 activity with a half maximum inhibitory concentration (IC_50_) of 1.50 nM (Figure S1, A). The binding affinity of MNL-4760 to ACE2 measured by MST was 702 nM (Figure S1, B). Consequently, MLN-4760 was used as a positive control in both assays to screen the compounds of the NCATS Pharmaceutical Collection and the anti-infectives library.

A total of 3149 compounds were screened in the ACE2 enzymatic assay in a 5-point dilution series with final compound concentrations ranging from 20 nM to 62 μM in 1536-well-plate format. The Z′-factor for the assay had an average value of 0.72 ± 0.04 and a signal-to-background ratio of 13.82 ± 2.73, indicating a robust assay performance. Compounds which showed ACE2 enzyme modulating activity in the primary screen were cherry-picked and retested in a 11-point dilution series ranging from 0.4 nM to 123.5 μM in duplicates. These compounds were tested in a counterscreen to check whether they interfere with the reporter fluorophore signal as well. Out of the 128 cherry-picked compounds 112 compounds were confirmed, where 110 of them were inhibitors of ACE2 enzyme and 2 activators.

The same small molecule libraries were screened for ACE2 binding by MST in 96-capillary-format at a single dose with final concentrations ranging from 39 to 392 μM depending on the highest available concentration of the compounds in the library due to their different solubility. Out of the 492 compounds (14.36%, Figure S1, C) identified as potential hits, 405 unique compounds were selected for the affinity screening at 7-point dilution series. The compounds were counterscreened with a fluorophore labeled His-peptide as well, to identify compounds interacting with the His-tag of the recombinant ACE2 instead of the target protein. The hit compounds identified from the affinity screen with dissociation constant (K_d_) values less than 30 μM and not active in the ACE2 enzymatic assay were re-tested in a second round of MST experiment. 116 compounds were validated as ACE2 putative allosteric binders.

### Virtual Screening

#### UNC Modeling

Model validation statistics for all evaluated models are reported in Supplementary Table 2. Models in bold were selected for final predictive use. According to standard metrics of model accuracy, model performance was fairly poor (correct classification rates close to 0.5). However, we placed emphasis on the positive predictive value (PPV) statistic, as our goal for validation was to determine the likelihood that any molecule that we nominate for experimental screening is truly active. We sought an increase in PPV over the selection strategy for the initial screening set (3.7% active rate).

Four of these models were selected to be used in consensus based on PPV and diversity of model type and descriptors. Our consensus model performance predicted about a 3-fold increase in the rate of active compounds. The strategy for nominating new compounds proceeded by first predicting every compound in the screening set with the MuDRA model and the gradient boosting model built on the RDKit descriptors. The simplex-based models take a significant amount of computation time to generate descriptors, so they were not included in the initial screening step. The gradient boosting model predicted a total of 20 actives, and the MuDRA model predicted 1015 actives, with no overlap between the two models.

This set of 1035 actives was further predicted by the simplex-based models for prioritization. 6 compounds were predicted active by both the MuDRA model and a simplex-based model and were considered the highest priority nominations. All 20 actives predicted by the gradient boosting model with RDKit descriptors were chosen as the next tier of nominations. To reach 256 nominated compounds, all remaining actives predicted by the MuDRA model were sorted by Euclidean distance to the nearest training set active compound in the RDKit descriptor space, and the top 230 were selected.

#### NCATS Modeling

Detailed model statistics on the training set and the test set are provided in Supplementary Table 3. All four models showed AUC values close or above 65% in both training set and test datasets. Taking a consensus between different descriptor combinations did not improve the model performance.

For the LBP modeling, we used the 110 active compounds; clustering based on pharmacophore-based similarity (cluster distance of 0.4, 0.6, 0.7, and 0.8), followed by generation of ligand-based hypotheses led to a total of 89 pharmacophore hypotheses (MFP and SFP). Taking the computational constraints into account, 24 pharmacophore models that hit the majority (>20%) of active vs. inactive were selected for virtual screening. All pharmacophore hypotheses generated in this study are presented in Supplementary Table 1. In general, the pharmacophoric sites such as hydrogen bond acceptor (HBA), hydrogen bond donor (HBD) and aromatic ring, were prudently characterized.

For the virtual screening, the complete collection of 138,729 compounds was tested against our SB models and ranked by the prediction score. This score takes a value between 0 and 1, and the higher the score, the higher the probability of the compound to be active. We filtered the compounds that have the prediction score greater than 0.60 from each descriptor combination. This led us to a subset of 660 compounds that were predicted to be active by at least two out of the four models. We also screened the 138,729 compounds using the LBP models shortlisted for the screening (Supplementary Table 1). This gave us 16,231 compounds, 58 of which were also in the list of 660 compounds (from SB). These 58 compounds were selected for the experimental validation. Owing to poor performance of our SB models we only picked another top 58 compounds from the SB approach (ranked according to their prediction score). The other 140 compounds were selected from our LBP models, according to their pharmacophore fit score (that hit at-least 3 pharmacophore models) and sorted by the Tanimoto similarity (based on Morgan fingerprints) to the nearest training set active compound. This gave us a list of 256 compounds for experimental validation.

### Post-modeling Screening and Follow-up Experiments

As described above, computational models were used to nominate a total of 512 compounds for experimental testing. Within the top 256 compounds, chosen from each institute, there were overlapping 11 compounds. Thus, we added additional 11 compounds from the UNC list (actives predicted by the MuDRA model and sorted by Euclidean distance to the nearest training set active compound in the RDKit descriptor space).

First, these molecules were tested for ACE2 binding by MST at a single concentration. Out of the 512 compounds, 130 (25.39%) were identified as binders and nine – as inconclusive (Figure 2). Next, the 139 potential hit compounds were measured by MST in dose-response at 7-point dilution series for ACE2 binding and in His-peptide counterscreen to test the binding specificity. These compounds were tested in ACE2 enzymatic assay as well. Sixty-eight compounds were identified as ACE2 binders with K_d_ values ranging from 6 nM to 562 μM, where only two of them had moderate enzyme inhibitory activity.

**Figure 1:**
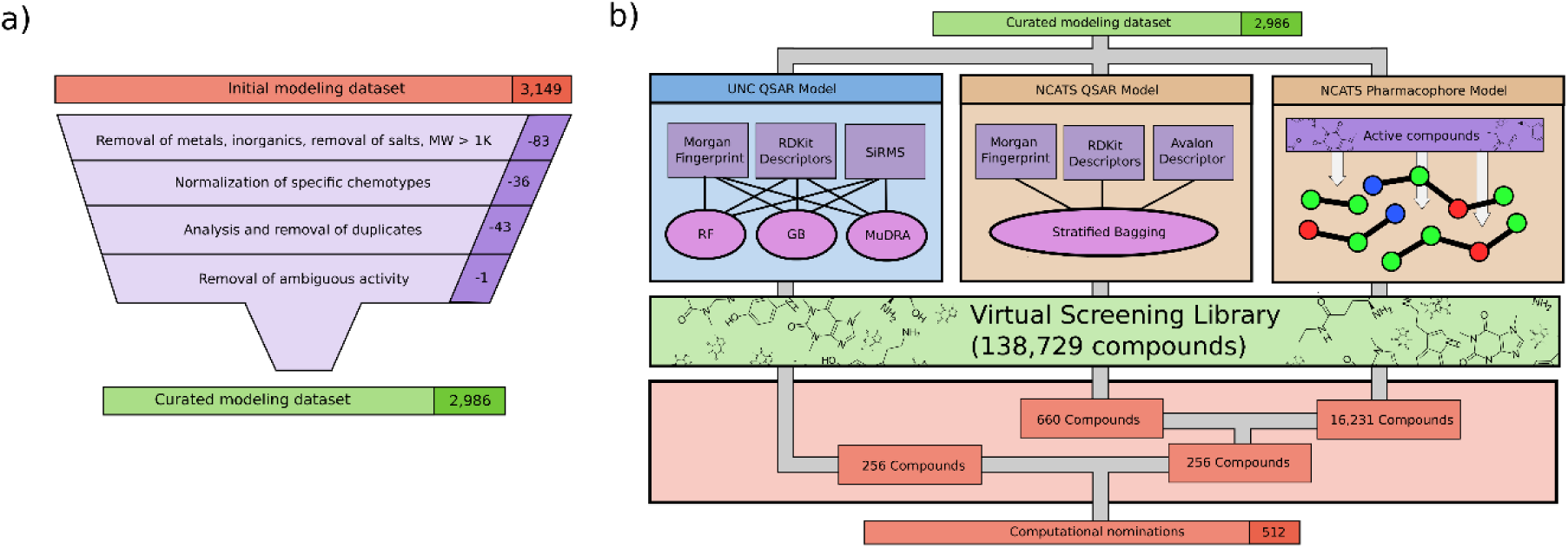
Computational workflow. a) Curation protocol. UNC curation protocol^34–36^ shown, and NCATS protocol is similar with minor deviations. b) Modeling workflow. In total, three model types were used to screen compounds to produce 512 nominations for further experimental validation.

**Figure 2.**
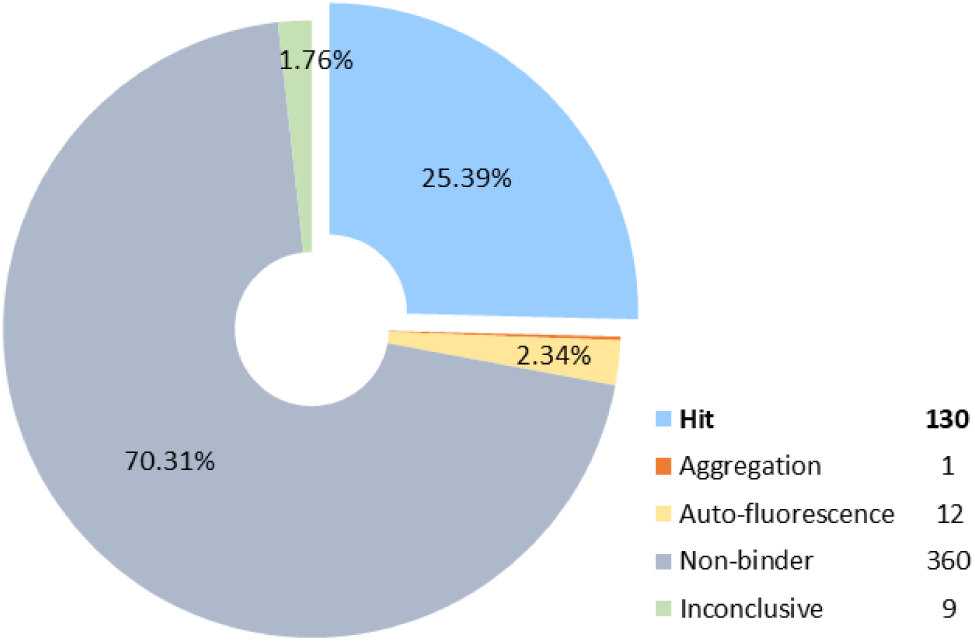
Experimental testing of the predicted molecules for ACE2 binding by MST at a single dose. Out of the tested 512 compounds, 130 (25.39%) were identified as hits, 360 as non-binders, 9 inconclusive. Twelve compounds were auto-fluorescent, and one compound caused aggregation.

To determine whether putative allosteric binders of ACE2 can affect SARS-CoV-2 infection, compounds with K_d_ below 10 μM and no enzyme inhibiting activity were tested in live SARS-CoV-2 Fluc assay (see the Material and Methods section for details). The assay indirectly monitors the ability of compounds to inhibit viral replication and infection through various molecular mechanisms, including direct inhibition of viral entry or enzymatic processes as well as acting on host pathways that modulate viral replication. The compounds were tested in the corresponding counterscreens for cytotoxicity and luciferase inhibitory activity.

Overall, 5 compounds showed a SARS-CoV-2 inhibiting activity with an efficacy greater than 70% and half maximal inhibitory concentration (IC_50_) values of 14 - 25 μM (Table 1). Of these compounds, only compound 1 showed a cytotoxic effect with IC_50_ = 24.8 μM (Figure S2, A), and compounds 3 and 5 had moderate luciferase inhibiting activity (Figure S2, B). The remaining compounds were inactive in the counterscreens and can be considered as true SARS-CoV-2 inhibitors.

**Table 1:**
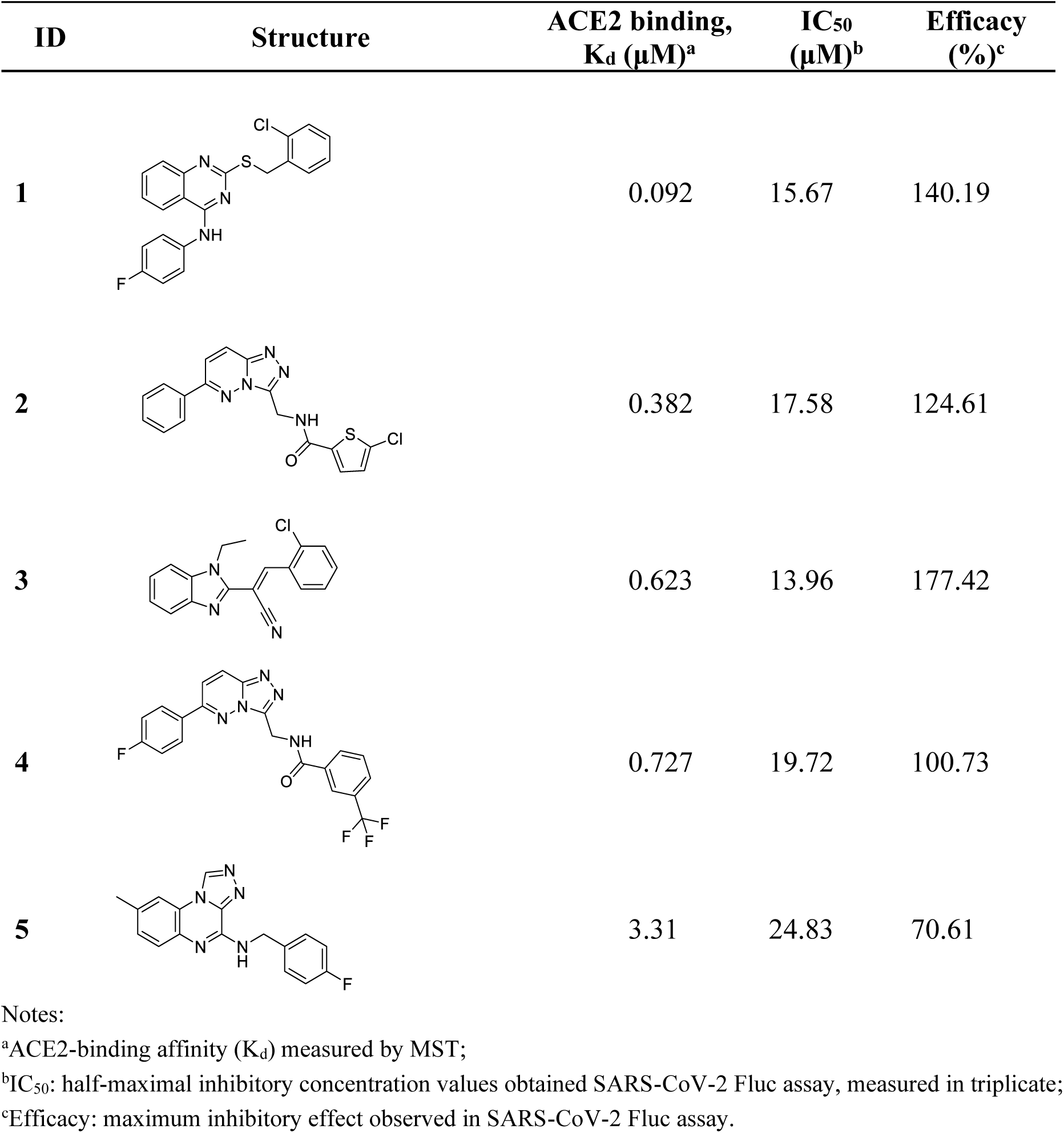
Compounds Identified as ACE2 Binders and Inhibitors of SARS-CoV-2

| ID | Structure | ACE2 binding,<br>K<sub>d</sub> (μM)<sup>a</sup> | IC<sub>50</sub><br>(μM)<sup>b</sup> | Efficacy<br>(%)<sup>c</sup> |
| --- | --- | --- | --- | --- |
| 1 |  | 0.092 | 15.67 | 140.19 |
| 2 |  | 0.382 | 17.58 | 124.61 |
| 3 |  | 0.623 | 13.96 | 177.42 |
| 4 |  | 0.727 | 19.72 | 100.73 |
| 5 |  | 3.31 | 24.83 | 70.61 |

Notes:
<sup>a</sup>ACE2-binding affinity (K<sub>d</sub>) measured by MST;
<sup>b</sup>IC<sub>50</sub>: half-maximal inhibitory concentration values obtained SARS-CoV-2 Fluc assay, measured in triplicate;
<sup>c</sup>Efficacy: maximum inhibitory effect observed in SARS-CoV-2 Fluc assay.

To summarize, we used a microscale thermophoresis assay (to measure compound binding to ACE2) and an enzymatic assay (to determine whether cleavage of a known peptide substrate of ACE2 was interrupted in the presence of these compounds) to discover putative allosteric binders of ACE2. We presumed that a compound that shows strong binding to ACE2 but little to no interruption of enzymatic activity is an allosteric binder. Using the results of the initial binding and enzymatic activity assays (3,246 compounds), we developed the QSAR models, which were used to nominate a set of 512 virtual hits. Experimental validation of these compounds demonstrated that 13% of them were allosteric ACE2 binders. This was a significant enrichment of hit rate over the prevalence of allosteric binders in the original assay, which was closer to 3%. This set of compounds, along with hits from the original assay, were then subjected to a further experimental analysis to determine their anti-SARS-CoV-2 activity. Thus, an assay for viral replication of a modified SARS-CoV-2 virus in human cells over-expressing ACE2 was applied to the set of presumed allosteric ACE2 binders, along with appropriate counterscreens to rule out assay artifacts. In total, five hit compounds reported here have significant binding to ACE2, little disruption of ACE2 enzymatic activity, and significantly reduce SARS-CoV-2 replication in human cells. To the best of our knowledge, this is the first successful attempt to discover antiviral agents against SARS-CoV-2 via ACE2 allosteric binding mechanism, and we encourage further exploration of these hits for use in treatment of COVID-19.

## Material and Methods

### Experimental

#### Microscale Thermophoresis Assay

Binding of the compounds to recombinant human ACE2 (Sino Biological, Cat #: 10108-H08H) was evaluated by microscale thermophoresis (MST). His-tagged ACE2 was labeled with REDtris-NTA 2nd generation dye (Nanotemper Technologies, Cat #: MO-L018) following manufacturer’s protocol and diluted in MST buffer (10 mM HEPES pH 7.4, 150 mM NaCl, 10 mM CaCl2, 0.01% Tween 20) to a final concentration of 3 nM For the single dose screen 200 nL of library compounds were pre-dispensed into the assay plate using Echo 650 series acoustic dispenser (Labcyte Inc.), mixed with 10 μL of labeled protein and incubated for 15 min at RT. For dose-response experiments, 100 nL of compounds in two-fold dilution series were transferred to 384-well compound plate (Greiner, Cat #: 784201-1B). MST traces were collected using Monolith NT.Automated (Nanotemper Technologies) unit and standard treated capillary chip (Nanotemper Technologies, Cat #: MO AK002) with following setting: 45% excitation power, medium MST power and MST periods of 3s/10s/1s. K_d_ values were calculated by fitting the change in normalized fluorescence signal of the thermograph using MO.Affinity analysis software.

To identify false-positive binders, which could interact with the fluorophore labeled His-tag instead of the target protein, all compounds tested in the dose-response experiments with ACE2 were counterscreened with poly-histidine control peptide (Nanotemper Technologies) under the same experimental conditions as with ACE2.

#### ACE2 Enzymatic Assay

ACE2 enzyme activity was monitored in a fluorometric assay. Briefly, 25 nL compounds were transferred to the 1536-well assay plate (Greiner, solid black medium-binding plates) using Echo 650 (Labcyte Inc.) acoustic dispenser. 3 μL/well of 0.27 nM ACE2 (0.2 nM final concentration) suspension in assay buffer (PBS, pH 7.4, 0.01% Tween-20) was dispensed into assay plate with Aurora Discovery BioRAPTR Dispenser (FRD; Beckton Dickenson) and incubated 15 min at room temperature (RT). One μL/well of 60 μM ACE2 substrate (AnaSpec, Cat #: AS-60757) was then added. The plate was centrifuged at 1000 rpm for 15 sec and the fluorescence was detected with the PHERAstar plate reader (BMG Labtech) equipped with Module 340/440 at t1=0 min and t2=15 min at RT. Data was normalized to enzyme activity in presence of DMSO, set as 0%, and in presence of 6.2 μM MLN-4760, set as -100% inhibition. The resulting percent of inhibition data were fitted to a sigmoidal dose response curve using four-parameter Hill equation.

#### Live SARS-CoV-2 Fluc Assay and Cytotoxicity Counterscreen

A live SARS-CoV-2 replication assay in A549-ACE2 host cells was used to measure the ability of compounds to perturb the replication of SARS-CoV-2. It employs an engineered SARS-CoV-2 WA-1 lineage virus that has an integrated firefly luciferase reporter (Fluc, provided by Pei-Yong Shi, UTMB) and A549-ACE2 cells (generously provided by Pei-Yong Shi, UTMB), an adenocarcinoma human alveolar basal epithelial cell line stably overexpressing human ACE2.

Briefly, 20 nL/well of compounds in DMSO were spotted into 1536-well assay plates (Aurora E8, black clear bottom, tissue culture treated plates) by acoustic dispensing. In parallel, 20 nL of DMSO were added to the first 4 columns of the plate, which serve as the no virus and neutral control wells. 4 μL of A549-ACE2 cells are dispensed to all wells, for a final density of 1,600 cells/well in DMEM media with 2% FBS. In addition, 1 μL of media (DMEM, 2% FBS) are dispensed to columns 1 and 2. Thereafter, 1 μl/well of SARS-CoV-2 (USA_WA1/2020) at multiplicity of infection (MOI) of 0.2 suspended in media were dispensed to columns 3-48. Assay plates were incubated for 48 h at 37°C, 5% CO_2_, 90% humidity. After incubation, 2 μL/well of One-Glo (Promega, Cat # E6120) detection reagent was added, and plates were incubated for 5 min at room temperature. Luminescence signal was measured on BMG PheraStar plate reader.

Raw data was normalized to the neutral control (cells infected with virus in presence of DMSO, set as 0%) and positive control (cells without virus added, set as -100%) for each plate. The resulting percent of inhibition data were fitted to a sigmoidal dose response curve using four-parameter Hill equation.

In parallel, the compounds were tested in a cytotoxicity counterscreen against the A549-ACE2 cell line. The assay was set up in the same way as in the Fluc assay omitting the addition of virus. A549-ACE2 with DMSO solvent served as the negative control, whereas media and DMSO (no cells) was the positive control. The plates were incubated for 48 hrs. at 37°C, and one volume of CellTiter-Glo assay reagent (Promega, Madison, WI), which assess viable cells (ATP content), was added using a BioRAPTR FRD (Beckman Coulter, Brea, CA). Cell viability was measured using a ViewLux μHTS Microplate Imager (PerkinElmer, Waltham, MA). The obtained luminescence signal was normalized against negative control (0% response) and positive control (−100% response).

#### Firefly Luciferase Counterscreen

To identify false-positive hits, which could reduce the Fluc signal due to the inhibition of the luciferase enzyme rather than perturbing the viral infection, compounds were tested in a biochemical assay with recombinant luciferase from *Photinus pyralis* (firefly). Briefly, 25 nL/well compounds or DMSO as vehicle control (columns 1-4) were acoustically transferred to white solid 1536-well-plate (Greiner, Cat #: 789175-F). 3 μL/well of 13.33 nM luciferase (10 nM final concentration) suspension in 50 mM Tris-acetate buffer, pH 7.6 was dispensed into assay plate with Aurora Discovery BioRAPTR Dispenser (FRD; Beckton Dickenson) and incubated 15 min at room temperature. 3 μL/well of buffer only were dispensed to columns 3-4 as no enzyme control. One μL/well of 40 μM D-luciferin in substrate buffer (50 mM Tris-acetate, pH 7.6, 10 mM Mg-acetate, 10 μM ATP, 0.01% Tween-20, 0.05% BSA) was then added. The plate was centrifuged at 1000 rpm for 15 sec and the luminescence was detected with the PHERAstar plate reader (BMG Labtech). Data was normalized to enzyme activity in presence of DMSO, set as 0%, and no enzyme control, set as -100% inhibition. The resulting percent of inhibition data were fitted to a sigmoidal dose response curve using four-parameter Hill equation.

### Computational

#### Data Curation

##### UNC protocol

All chemical structures and correspondent activity information were analyzed and prepared according to data curation protocols proposed by Fourches et al.^34–36^ All steps of data curation were performed in KNIME^37^ integrated with ChemAxon Standardizer^38^. In summary, specific chemotypes were normalized and explicit hydrogens were added. If present, polymers, salts, organometallic compounds, and mixtures were removed. Furthermore, we performed the analysis and exclusion of duplicates. The following criteria were applied for exclusion of duplicates: (i) if the reported activity of the duplicates were the same (i.e., in concordance), only one entry was kept in the dataset; (ii) if duplicates presented discordance in biological activity, both entries were excluded. 1 compound with ambiguous activity data was removed. In total, 163 compounds were removed in the curation process, 6 of which were actives.

NCATS followed similar protocols using KNIME^37,39^ and the Atkinson standardization protocol, described at https://github.com/flatkinson/standardiser.^40^

#### Molecular Descriptors

##### UNC protocol

Descriptors were chosen to cover a reasonable amount of descriptor types (whole molecule descriptors, fragment descriptors, topological descriptors) while maximizing ease-of-use. A total of seven different descriptors were generated for the dataset. Six of these (RDKit whole-molecule descriptors, Morgan fingerprint, HashedAtomPairFingerprint, HashedTopologicalTorsionFingerprint, and MACCS Keys) were generated using the RDKit^41^ Python package.

The seventh descriptor type, Simplex Representation of Molecular Structures (SiRMS), was calculated using the Hit QSAR software.^42^ At the 2D level, the connectivity of atoms in a simplex, the atom type, and bond nature (single, double, triple, aromatic) were considered^43^. Bonded and nonbonded 2D simplexes were used. In addition to element and atom type, physicochemical characteristics of atoms, such as partial charge, lipophilicity, refraction, and the atom’s ability to be a hydrogen-bond donor/acceptor, were used for atom differentiation in the simplexes. For the atom characteristics with real values (charge, lipophilicity, and refraction), a binning procedure was used to define discrete groups: (i) partial charge A ≤−0.05 < B ≤ 0<C ≤ 0.05 < D, (ii) lipophilicity A ≤−0.5 < B ≤ 0<C ≤ 0.5 < D, and (iii) refraction A ≤ 1.5 < B ≤ 3<C ≤ 8<D. For hydrogen-bond characteristics, the atoms were also divided into three groups: A (acceptor of hydrogen in H-bond); D (donor of hydrogen in H-bond); and I (indifferent atom, i.e., atom that does not form H-bonds)^44^.

##### NCATS protocol

We employed three different sets of descriptors: physicochemical descriptors (RDKit), Morgan fingerprints (1024 bits) and Avalon fingerprints (1024 bits), calculated using the RDKit toolkit^41^. As consensus modeling approaches have been reported to outperform simple QSAR models^45–49^, we also performed the consensus of descriptors (RDKit, Morgan and Avalon).

#### Model Building

##### UNC QSAR protocol

The models were developed using best practices as described in Cherkasov et al.^50^ For models not based on decision trees, the descriptor matrix was normalized to prevent undesired weighting of certain descriptor dimensions. Descriptor dimensions with low variance (less than 0.0001) were eliminated due to being non-informative. Models were generated for most architecture-descriptor pairs. The MuDRA^51^ algorithm requires multiple descriptor spaces, so it was handled differently than the rest. The four descriptor types used for the MuDRA model were the Morgan fingerprint, HashedAtomPairFingerprint, HashedTopologicalTorsionFingerprint, and MACCS Keys.

##### NCATS QSAR protocol

In order to overcome the problem of data imbalance, we used bagging with stratified under-sampling^52^. This method has proven to be among the best performing methods for dealing with imbalanced datasets^53^. Stratified bagging (SB) is a machine learning technique that is based on an ensemble of models developed using multiple training datasets sampled from the original training set. It uses minority class samples to create the training set of positive samples using a traditional bagging approach (resampling with replacement) and after that randomly selects the same number of samples from the majority class. Thus, the total bagging training set size was double the number of the minority class molecules. Several models are then built, and predictions averaged in order to produce a final ensemble model output. Because of random sampling, about 37% of the molecules are not selected and left out in each run. These samples create the “out-of-the-bag” sets, which are used for testing the performance of the final model^53^. Although a small set of samples are selected each time, the majority of molecules contributed to the overall bagging procedure since the datasets were generated randomly. Random Forest (with default parameters) was used as a base-classifier. The number of trees was arbitrarily set to 100 (default), since it has been shown that the optimal number of trees is usually 64-128, while further increasing the number of trees does not necessarily improve the model’s performance.

##### Pharmacophore-based modeling

In addition to QSAR, we also performed ligand-based pharmacophore modeling (LBP). A pharmacophore describes the spatial arrangement of essential interactions of a drug with its respective receptor binding site. It is a well-established method virtual screening (VS) in the early drug discovery process. In this study, the generation of LBP models, their subsequent refinement, and VS were performed with LigandScout 4.4 Advanced, available by Inte:Ligand GmbH. The conformational libraries for both pharmacophore modeling and the VS process were created with i:Con^54^ (max. 200 conformations per compound), a conformer generator implemented in LigandScout.

To design the LBP models, the actives (from the training set) were clustered based on pharmacophore-based similarity (cluster distance 0.4, 0.6, 0.7, and 0.8, respectively). For each of the clusters obtained from different cluster distance thresholds, merged-features pharmacophore (MFP) and shared-features pharmacophore (SFP) models were generated that incorporate the features of selected compounds per cluster.^55^ A good pharmacophore model should not only be able to estimate the activity of active compounds, but also have the ability to identify the active molecules from a database containing a large number of inactive compounds. To select the best models for screening, we applied these models on our complete dataset (training and test set combined) and calculated the percentage of active and inactive that hit these pharmacophore models. The models that hit 20% more active compounds versus inactive compounds were selected for the final virtual screening. The screening was performed using iscreen module, with default settings with the maximum number of omitted features set to 2.

#### Model Validation

##### UNC protocol

Models were evaluated using five-fold external cross-validation. The dataset was split randomly into five partitions. Each model was trained on four of the five partitions and tested on the fifth. This process was repeated five times so each partition was used once as a test set. Reported model statistics are an average of performance across each test set.

##### NCATS protocol

From each class, 80 % of the data was randomly selected and used as a training set. The remaining 20 % of compounds were considered as the external validation set. For stratified Bagging, since multiple training datasets were generated by selecting the molecules with replacement from training set in a random fashion, this leaves out about 37% of the instances in each run. Therefore, these molecules that constitute the ‘out-of-the-bag’ sets are later used for testing the performance of the final model.

## Data availability

All the data, models, and results produced/developed in this study are available at Chembench^56^(https://chembench.mml.unc.edu/).

## Author contribution

Conceptualization, ENM, AT, BB, and AVZ; Methodology, JEH, SJ, RE, ENM, AT; Validation, JEH, CCMF; Assay conceptualization AS; Experimental validation, BB, RE; qHTS analysis and selection of chemotypes GR, DCT; Formal Analysis, LC, HG. Investigation, TB, ZLS, ENM; Data Curation, CCMF, TB, ZLS, SJ; Writing – Original Draft, JEH and SJ, Writing – Review & Editing, all authors; Visualization, JEH and BB.

## Acknowledgement

Authors from UNC-Chapel Hill were supported by National Institutes of Health (Grants OT2TR002514 and R01GM140154). JEH was supported by National Institutes of Health Training Grant (T32GM008570). This research was supported by the Intramural Research Program of the National Center for Advancing Translational Sciences (NCATS), National Institutes of Health (NIH).

