## Supplementary Information for "Allosteric binders of ACE2 are promising anti-SARS-CoV-2 agents"

Supplementary Table 1: Ligand-based pharmacophore models generated for screening (LBP).

| Pharmacophore | pharmacophore type | cluster distance |
|---|---|---|
| H-HBA-HBA-HBD-HBD-NI-XV44 | MFP | 0.4 |
| AR-AR-AR-H-HBA-HBA-HBA-HBD-HBD-HBD-XV46 | MFP | 0.4 |
| AR-AR-H-H-HBA-HBA-HBA-HBA-HBD-XV42 | MFP | 0.4 |
| H-HBA-HBA-HBA-HBA-HBD-HBD-XV34 | MFP | 0.4 |
| HBA-HBA-HBA-HBA-HBA-HBD-HBD-HBD-XV37 | MFP | 0.4 |
| AR-AR-AR-H-HBA-HBA-HBA-HBD-HBD-HBD-XV46 | MFP | 0.6 |
| H-HBA-HBA-HBA-NI-XV42 | MFP | 0.6 |
| HBA-HBA-HBA-XV27 | MFP | 0.6 |
| AR-AR-H-H-HBA-HBA-HBA-HBD-XV35 | MFP | 0.6 |
| AR-AR-AR-HBA-HBA-HBA-HBA-HBA-HBD-XV47 | MFP | 0.6 |
| AR-H-HBA-HBA-HBA-HBA-HBD-HBD-HBD-XV41 | MFP | 0.6 |
| AR-HBA-HBA-HBA-HBD-XV37 | MFP | 0.6 |

| | | |
|---|---|---|
| HBA-HBA-HBA-HBA-HBA-HBA-HBD-PI-XV40 | MFP | 0.6 |
| AR-AR-H-HBD-XV29 | MFP | 0.6 |
| HBA-HBD-HBD-HBD-XV27 | MFP | 0.6 |
| HBA-HBA-HBD-XV26 | MFP | 0.6 |
| HBA-HBA-HBA-XV19 | MFP | 0.6 |
| H-HBA-HBA-XV21 | MFP | 0.6 |
| AR-HBA-HBA-HBA-XV38 | MFP | 0.6 |
| AR-AR-H-HBA-HBA-HBD-HBD-NI-XV37 | MFP | 0.6 |
| AR-H-HBA-HBA-HBD-HBD-XV29 | MFP | 0.7 |
| AR-AR-H-HBD-XV30 | MFP | 0.7 |
| AR-H-HBA-HBA-HBA-HBD-NI-XV36 | MFP | 0.7 |
| AR-HBA-HBA-HBD-XV21 | MFP | 0.7 |
| AR-H-HBA-HBA-XV13 | MFP | 0.7 |
| HBA-HBA-HBD-HBD-XV21 | MFP | 0.7 |
| HBA-HBA-HBD-XV18 | MFP | 0.7 |
| AR-H-HBA-HBA-HBA-HBD-XV34 | MFP | 0.7 |
| HBA-HBA-HBA-XV28 | MFP | 0.7 |
| H-H-HBA-HBA-NI-XV33 | MFP | 0.7 |
| HBA-HBA-HBA-HBD-XV22 | MFP | 0.7 |
| HBA-HBA-HBD-HBD-HBD-HBD-XV39 | MFP | 0.7 |
| H-HBA-HBA-HBA-HBD-XV31 | MFP | 0.7 |
| AR-H-HBA-HBD-XV24 | MFP | 0.8 |
| HBA-HBA-HBA-XV18 | MFP | 0.8 |
| HBA-HBA-HBA-HBD-XV1 | MFP | 0.8 |
| H-HBA-HBA-XV31 | SFP | 0.4 |
| AR-AR-AR-H-HBA-HBA-HBA-XV34 | SFP | 0.4 |
| AR-AR-AR-H-XV17 | SFP | 0.4 |
| AR-HBA-HBA-XV25 | SFP | 0.4 |
| AR-AR-HBA-HBA-HBD-XV24 | SFP | 0.4 |
| H-HBA-HBA-HBA-HBD-XV32 | SFP | 0.4 |
| AR-AR-H-H-H-XBD-XV31 | SFP | 0.4 |
| HBA-HBA-HBA-HBA-HBD-HBD-XV35 | SFP | 0.4 |
| AR-AR-AR-H-HBA-HBA-HBA-XV34 | SFP | 0.6 |
| H-HBA-HBA-NI-XV36 | SFP | 0.6 |
| AR-HBA-HBA-XV25 | SFP | 0.6 |
| AR-H-H-HBA-XV30 | SFP | 0.6 |
| AR-H-HBA-HBA-HBD-XV33 | SFP | 0.6 |
| AR-H-HBA-HBA-HBD-HBD-XV36 | SFP | 0.6 |

| | | |
|---|---|---|
| AR-H-HBA-HBD-XV32 | SFP | 0.6 |
| HBA-HBA-HBA-HBD-PI-XV35 | SFP | 0.6 |
| HBA-HBA-HBA-NI-XV24 | SFP | 0.6 |
| AR-AR-H-XV13 | SFP | 0.6 |
| AR-H-HBA-XV17 | SFP | 0.6 |
| H-HBA-HBA-HBD-XV29 | SFP | 0.6 |
| HBA-HBA-HBD-XV21 | SFP | 0.6 |
| AR-H-HBA-XV22 | SFP | 0.6 |
| AR-HBA-HBA-XV22 | SFP | 0.6 |
| HBA-HBA-HBD-XV20 | SFP | 0.6 |
| HBA-HBA-HBA-XV16 | SFP | 0.6 |
| AR-HBA-HBA-XV28 | SFP | 0.6 |
| AR-AR-H-HBA-XV27 | SFP | 0.6 |
| H-HBD-HBD-XV30 | SFP | 0.7 |
| AR-AR-H-XV19 | SFP | 0.7 |
| HBA-HBA-HBD-NI-XV17 | SFP | 0.7 |
| H-HBA-HBA-HBD-HBD-XV28 | SFP | 0.7 |
| AR-H-H-HBA-XV31 | SFP | 0.7 |
| HBA-HBA-HBD-XV20 | SFP | 0.7 |
| HBA-HBA-HBD-XV21 | SFP | 0.7 |
| H-HBA-HBA-HBD-XV26 | SFP | 0.7 |
| AR-H-HBA-HBD-XV28 | SFP | 0.7 |
| HBA-HBD-HBD-XV27 | SFP | 0.7 |
| H-HBA-HBA-XV22 | SFP | 0.7 |
| HBA-HBA-NI-XV18 | SFP | 0.7 |
| H-HBA-HBA-XV28 | SFP | 0.7 |
| HBA-HBA-HBA-HBA-HBD-XV37 | SFP | 0.7 |
| H-HBA-HBA-XV29 | SFP | 0.7 |
| H-HBA-HBA-XV29 | SFP | 0.7 |
| HBA-HBA-HBA-XV23 | SFP | 0.8 |
| HBA-HBA-HBD-XV16 | SFP | 0.8 |
| AR-HBA-HBD-XV8 | SFP | 0.8 |
| H-H-HBA-XV24 | SFP | 0.8 |
| AR-HBA-HBA-XV21 | SFP | 0.8 |
| AR-HBA-HBA-XV11 | SFP | 0.8 |
| H-HBA-HBA-XV18 | SFP | 0.8 |
| H-H-HBA-XV39 | SFP | 0.8 |
| AR-AR-H-XV17 | SFP | 0.8 |
| HBA-HBA-HBD-XV4 | SFP | 0.8 |

| | |
|---|---|
| MEP | merged-features pharmacophore |
| SFP | shared-features pharmacophore |
| H | Hydrophobic |
| HBA | H-Bond Acceptor |
| HBD | H-Bond Donor |
| PI | Positive Ionizable |
| NI | Negative Ionizable |
| AR | Aromatic Ring |
| XV | Exclusion Volume |
| (green box) | pharmacophore models that hit the majority (>20%) of active vs. inactive |

Supplementary Table 2: Validation statistics for UNC models. Reported statistics are an average across each of five test folds. Final selected models are shown in **bold.** Acronyms: Correct Classification Rate (CCR), Positive Predictive Value (PPV), Negative Predictive Value (NPV).

| **Model Type** | **Descriptor** | **CCR** | **PPV** | **NPV** |
|---|---|---|---|---|
| **Neural Network** | **Simplex** | **0.514** | **0.096** | **0.964** |
| Neural Network | RDKit | 0.500 | 0 | 0.963 |
| Neural Network | Morgan | 0.500 | 0 | 0.963 |
| **Gradient Boosting** | **Simplex** | **0.516** | **0.123** | **0.965** |
| **Gradient Boosting** | **RDKit** | **0.511** | **0.102** | **0.964** |
| Gradient Boosting | Morgan | 0.495 | 0 | 0.963 |
| **MuDRA** | **MuDRA** | **0.522** | **0.129** | **0.965** |

Supplementary Table 3: Model performance of NCATS Stratified bagging models on the training and test dataset. Acronyms: Matthews correlation coefficient (MCC), Positive Predictive Value (PPV), Negative Predictive Value (NPV), Area under the receiver operating characteristic curve (AUC).

| **Training set** | | | | | | | | | |
|---|---|---|---|---|---|---|---|---|---|
| **Descriptor** | **Sensitivity** | **Specificity** | **Accuracy** | **Balanced Accuracy** | **MCC** | **Precision/PPV** | **NPV** | **Recall** | **AUC** |
| **RDKit** | 0.48 | 0.78 | 0.76 | 0.63 | 0.11 | 0.07 | 0.98 | 0.48 | 0.69 |
| **Morgan** | 0.30 | 0.87 | 0.85 | 0.58 | 0.09 | 0.08 | 0.97 | 0.30 | 0.69 |
| **Avalon** | 0.46 | 0.82 | 0.81 | 0.64 | 0.13 | 0.09 | 0.98 | 0.46 | 0.68 |
| **Consensus** | 0.38 | 0.85 | 0.83 | 0.61 | 0.12 | 0.09 | 0.97 | 0.38 | 0.72 |
| **Test set** | | | | | | | | | |
| **RDKit** | 0.41 | 0.83 | 0.81 | 0.62 | 0.11 | 0.08 | 0.97 | 0.41 | 0.62 |
| **Morgan** | 0.32 | 0.92 | 0.90 | 0.62 | 0.16 | 0.14 | 0.97 | 0.32 | 0.70 |
| **Avalon** | 0.27 | 0.87 | 0.85 | 0.57 | 0.08 | 0.07 | 0.97 | 0.27 | 0.57 |
| **Consensus** | 0.32 | 0.88 | 0.86 | 0.60 | 0.11 | 0.09 | 0.97 | 0.32 | 0.64 |

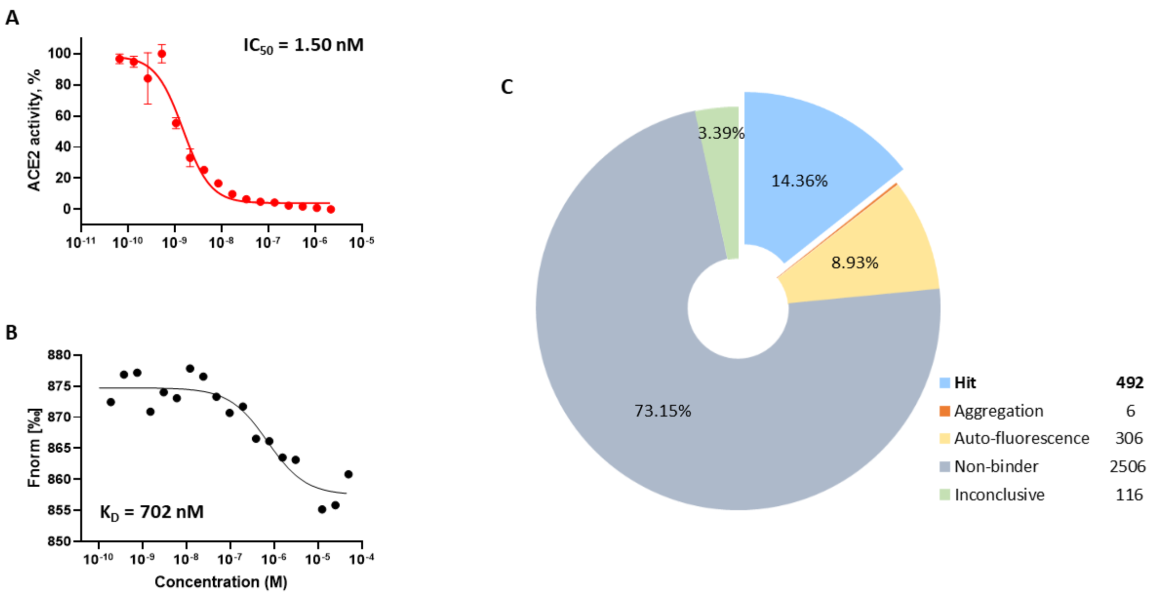


Figure S1

Supplementary Figure 1: Known ACE2 inhibitor MLN-4760 was tested in ACE2 screening assays to use as the control compound during screening. (A) Dose-response curve of MLN-4760 in ACE2 enzymatic assay with $IC_{50}$ of 1.5 nM, (B) MNL-4760 binding to ACE2 measured by MST. (C) Summary of the small molecule library screening for ACE2 binders using MST grouped by compound categories with total number of compounds in each group.

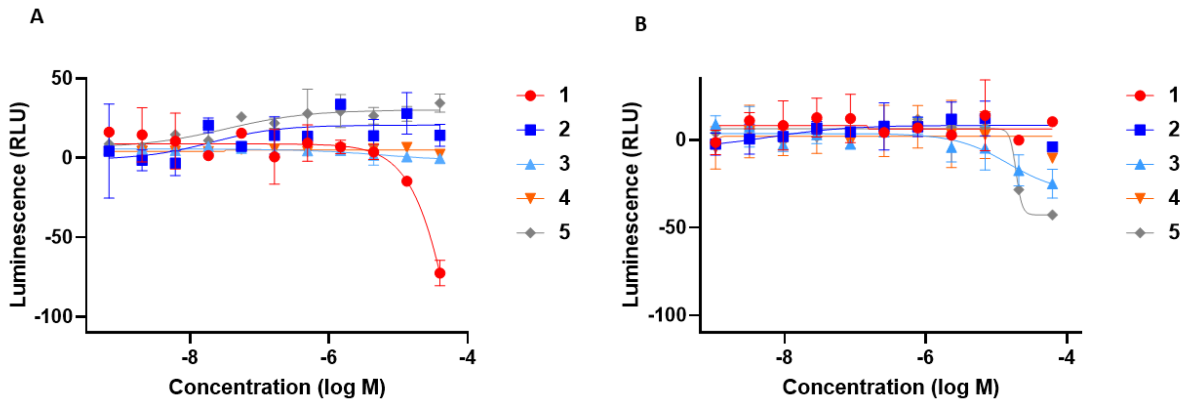


Supplementary Figure 2: Performance of the five ACE2 binders which showed inhibitory activity in the SARS-CoV-2 Fluc assay in the counterscreens: (A) cytotoxicity assay in A549-ACE2 cells, (B) firefly luciferase enzymatic assay.
